# Uncertainty in joint Ancestral State Reconstruction: Improving accuracy and biological interpretability of ancestral state prediction

**DOI:** 10.1101/2025.10.09.681367

**Authors:** James D. Boyko, Kyle J. Gontjes, Evan S. Snitkin, Stephen A. Smith

## Abstract

Ancestral state reconstruction (ASR) is a foundational tool in comparative biology, offering insights into the evolutionary history of lineages. With each new evolutionary model, our ability to estimate ancestral states has improved alongside the increased biological realism of these models. However, the field has primarily relied on reconstructions that focus on individual nodes, known as marginal reconstructions. This framework is analytically tractable but may not accurately represent what biologists want in inference, as evolution is dependent, and phenotypic transitions deeper in time can lead to consistent changes later. We argue that evolutionary history is better represented by joint reconstructions, which estimate the full sequence of states across nodes. Traditionally, joint reconstruction algorithms only estimated the single most likely sequence, but here we develop novel algorithms to estimate all relevant ancestral histories efficiently and provide tools to quantify the uncertainty of joint ASR. Furthermore, through simulations and an empirical case study, we demonstrate that joint reconstructions have higher accuracy than their marginal counterparts, and that the uncertainty surrounding the best joint reconstruction can be biologically meaningful and summarized using novel clustering algorithms. We apply our methods to epidemic multidrug-resistant *Klebsiella pneumoniae* and find that the evolution of antibiotic resistance is not a single narrative but a series of competing histories. Each of these histories exhibits distinct phenotype-genotype transitions that traditional approaches would struggle to identify, yet have critical implications for predicting resistance evolution.

## Introduction

Ancestral state reconstruction (ASR) is central to many, if not most, comparative studies in evolution. Typically based on observed state data ranging from genomic sequences to geographic areas to phenotypic descriptions (1–3), a fitted model is used to infer the unobserved states of ancestral lineages. This practice has become a valuable tool for comparative biologists because it provides context for the order of evolutionary events, often representing the only window into the unobserved history of life. ASR can also generate testable hypotheses by inferring ancestral states, which can subsequently be synthesized in the laboratory (e.g., 4). For example, reconstructed ancestral steroid receptors have been used to determine when specificity for different hormones evolved and demonstrated that several functional properties had been lost in descendant lineages (5). Even when reconstruction is not the primary goal, ASR approaches are an essential intermediate step of many other comparative analyses (2, 3, 6).

The increasing integration of phylogenetic perspectives in medical sciences has underscored the value of ancestral state reconstruction (2, 7). Phylogenetic comparative approaches have been successfully applied to model cancer prevalence and susceptibility across vertebrates (8), identify ancient enzymatic functions so that they could be compared to modern metabolic disorders (9, 10), and reconstruct ancestral states of virulence factors and antibiotic resistance genes (11–13). Reconstructing ancestral states has proven particularly powerful for understanding the evolutionary history of antibiotic resistance. By distinguishing between *de novo* evolution within hosts and the transmission of resistant strains, ancestral reconstructions have improved our understanding of how antibiotic resistance emerges and spreads (14–16). Despite its increasing application, little attention is given to whether the methodological foundations of ASR align with the biological questions that researchers seek to answer.

For historical and computational reasons, the field has primarily relied on specifying ancestral states based on marginal probabilities (2, 3, 17). Although ASR initially relied on parsimony algorithms for its estimates (e.g., 18), it quickly expanded to include maximum likelihood (19) and Bayesian (20) methods. With these likelihood-based methods, a parametric model describes node states as a weighted average of all the different scenarios given that the node is in a particular state (marginal probabilities). Essentially, for a given node, you are asking the probability that a state was present, considering every possible historical scenario (Figure 1). However, for at least some of the common questions asked by biologists, marginal probabilities focused on individual nodes may not be the best representation of ancestral states (21–23). In contrast, joint reconstruction methods aim to estimate the most probable configuration of states across all nodes simultaneously (Figure 1). This approach considers the likelihood of any given history and thus can provide biologists the most likely scenario when considering the entire evolutionary history of a clade (24). While marginal reconstructions provide uncertainty in the form of probabilities for a given state, algorithms for estimating the joint reconstruction have focused primarily on finding only the single best reconstruction (24). Even if the joint reconstruction aligns more closely with the types of questions in which biologists are interested, the potentially misleading uncertainty of the marginal reconstruction are still better than a joint reconstruction without uncertainty (25). What is needed is a method for estimating and representing the uncertainty around a joint estimation, as well as the tools for describing and utilizing that uncertainty in an empirical setting.

**Figure 1.**
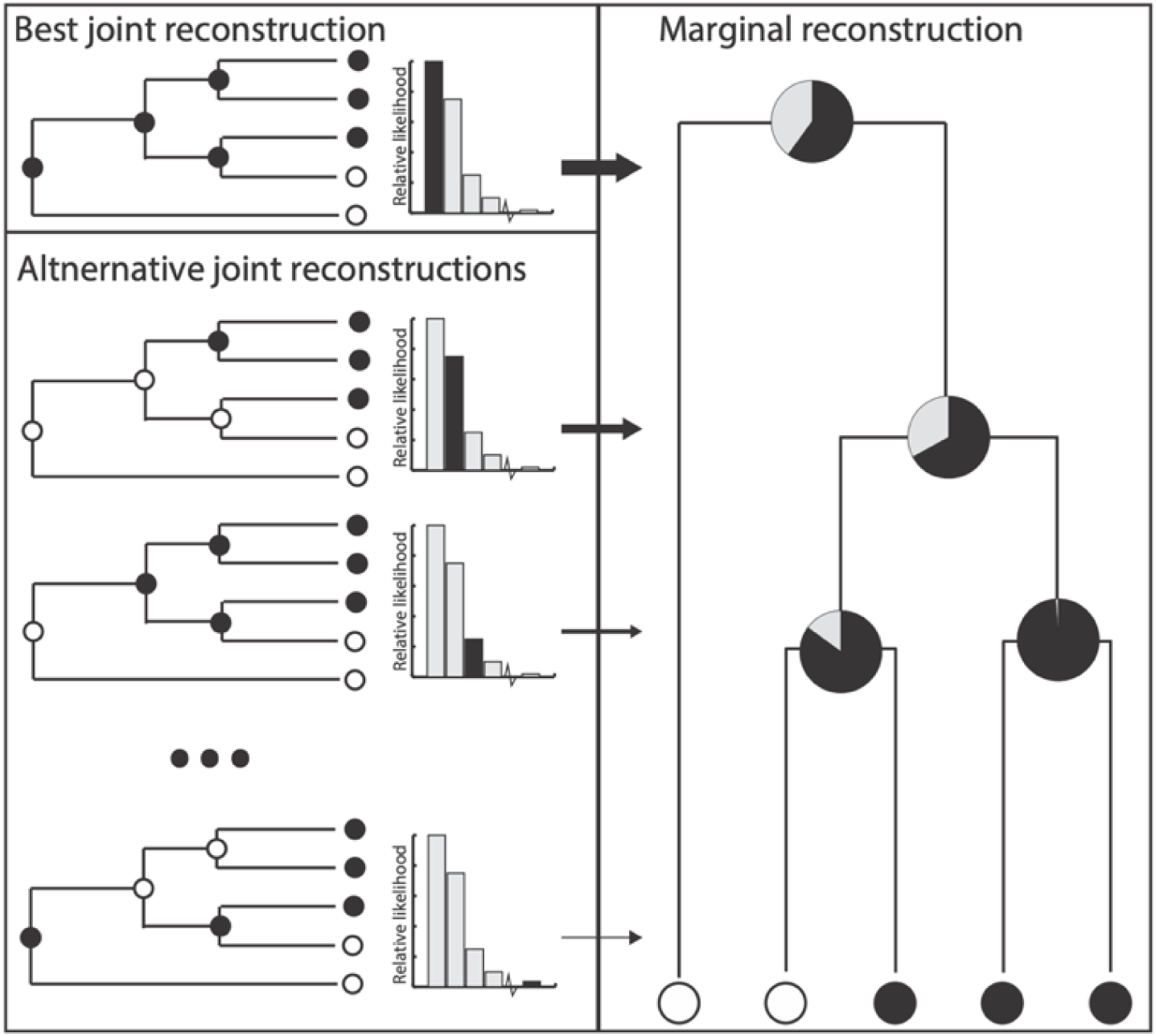
The relationship between the joint reconstruction and marginal reconstruction. The marginal reconstruction can be considered as a weighted average of all possible joint reconstructions. In doing so, the marginal effectively integrates over all alternative possibilities and presents a picture that is independent at each node. In contrast, the joint reconstruction represents a specific sequence of events, but current algorithms only return the single best reconstruction. This means that there are several other reconstructions which are currently ignored. Even though these alternatives may have a lower relative likelihood, they can represent a plausible evolutionary history that is quite different from the single most likely possibility. These alternatives can change inferences concerning when and how many times a phenotype emerges, as well as influencing downstream analyses.

We present two algorithms for estimating uncertainty for phenotypic and genomic information. Additionally, we develop a suite of methods for reconstruction clustering that facilitate the discovery of “major axes of ancestral reconstruction variation.” Our phylogenetically aware clustering approach allows empiricists to examine a smaller and more representative set of reconstructions than either the marginal or single joint estimates can provide. Each of these representative reconstructions can then be used as testable alternative hypotheses in downstream analyses. To test our methodology, we first compare the accuracy of marginal and joint ancestral estimation. Next, we evaluate the potential improvement by incorporating uncertainty around both marginal and joint reconstructions, examining whether our uncertainty intervals capture the true sequence of ancestral states. Finally, we develop several auxiliary tools necessary to apply these algorithms in an empirical setting and apply our novel framework to study the evolution of resistance to last-resort antibiotics in carbapenem-resistant *Klebsiella pnuemoniae*.

## Results

### Joint reconstruction outperforms marginal

Our first simulation study is designed to examine whether the single best marginal or joint ancestral estimate is more effective in identifying the true generating history across several simulation scenarios (see Methods). Reconstructions are generated using the same fitted model for each dataset so that only the reconstruction method differs. To test for systematic differences, we performed McNemar’s tests across all simulation conditions (26). The overall test revealed significant differences between the methods (χ² = 88.804, p < 2.2e-16), with the joint reconstruction consistently achieving higher accuracy rates than marginal reconstruction. We compared their accuracy across 1,000 simulated datasets for each parameter combination, finding that both methods exhibited moderate to low accuracy depending on the simulation conditions, but the joint method consistently outperformed marginal estimates (Figure 2).

**Figure 2.**
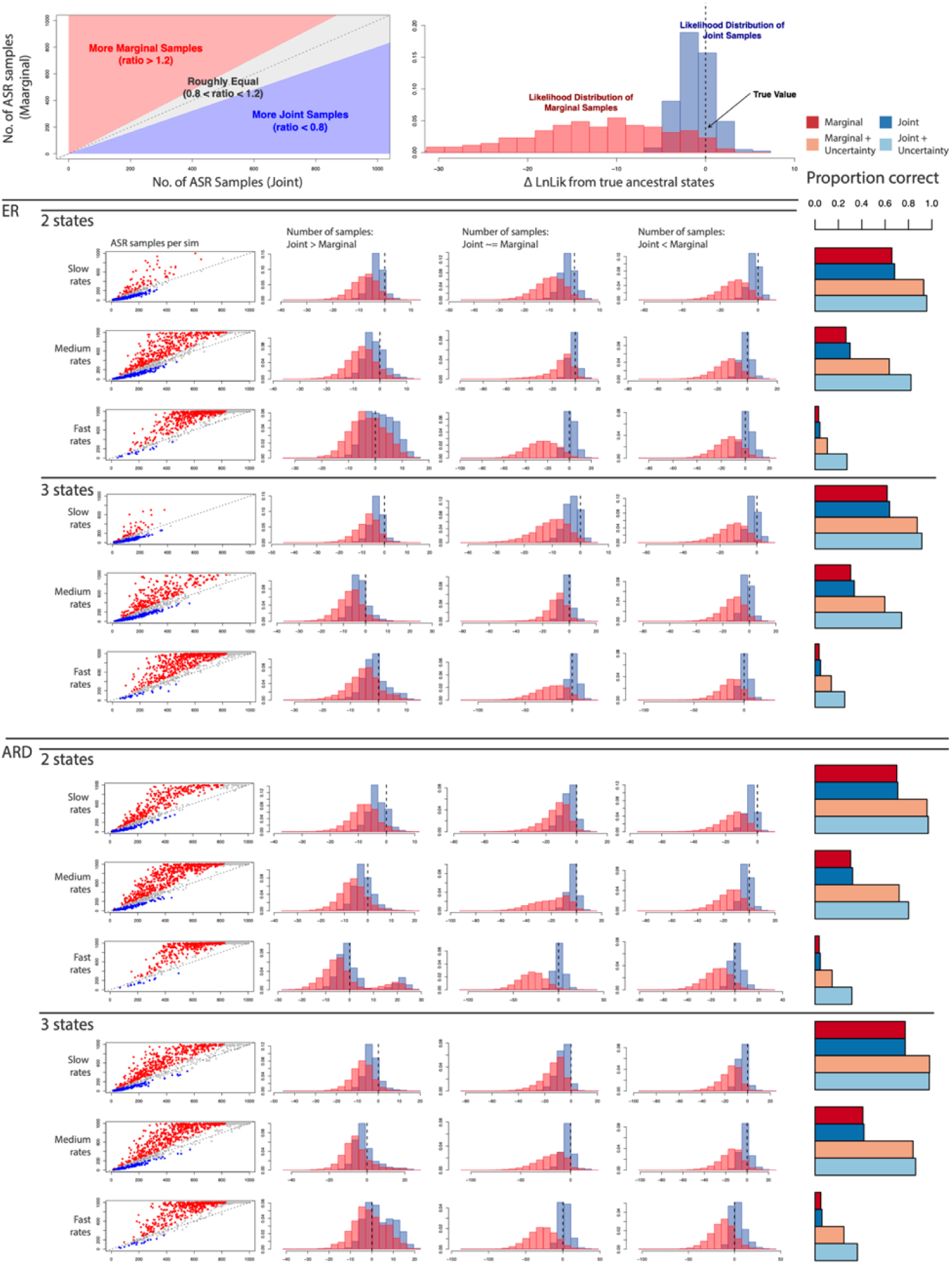
Comparison of joint and marginal uncertainty quantification methods. Visual key showing two components: (left) Sampling comparison where the x-axis represents the number of unique joint samples and the y-axis represents the number of marginal samples per simulation. Blue region indicates simulations where joint sampling produced more unique reconstructions, red region where marginal sampling produced more, and gray region where sampling was approximately equal. (right) Likelihood distribution comparison showing the difference between log-likelihood of the generating (true) reconstruction and reconstructions generated by each method. Red distribution represents marginal samples, and blue distribution represents joint reconstructions. The x-axis shows the difference in log-likelihood from the truth, with negative values indicating worse likelihood than the truth and positive values indicating better likelihood than the truth. Simulation results were organized into four scenarios: ER model with 2 states, ER model with 3 states, ARD model with 2 states, and ARD model with 3 states. The first column shows number of samples per simulation, color-coded according to panel a. Subsequent columns show likelihood distributions for each sampling regime, demonstrating that joint reconstructions (blue) generally center on the truth with narrower distributions across all simulation conditions. Bar charts corresponding to each simulation scenario were constructed to show the proportion of simulations in which the best marginal reconstruction, best joint reconstruction, marginal with uncertainty intervals, and joint with uncertainty intervals contained the true ancestral states. Results consistently show joint methods outperforming marginal methods, particularly at medium and high evolutionary rates.

We evaluated method performance under different model complexities by testing both an equal-rates (ER) model, where all transition rates between states were constrained to be equal, and an all-rates-different (ARD) model, where each transition rate was estimated independently. Under the correctly specified ER model (matching our simulation conditions), at low rates (0.25), joint and marginal methods showed identical accuracy for three-state characters (both 77.2%, McNemar’s χ² = 0, p = 1.0) and similar performance for two-state characters (joint: 70.8%, marginal: 70.1%, McNemar’s χ² = 1.89, p = 0.169). At medium rates (0.5), accuracy declined substantially for both methods, with joint reconstruction maintaining a slight edge (two-state: 32.2% vs 30.5%, McNemar’s χ² = 7.31, p = 0.007; three-state: 41.9% vs 40.9%, McNemar’s χ² = 2.13, p = 0.144). Finally, at high rates (1.0), both methods performed poorly, with accuracy dropping below 6% (two-state: 4.4% vs 3.6%, McNemar’s χ² = 3.5, p = 0.061; three-state: 5.9% vs 5.0%, McNemar’s χ² = 4.27, p = 0.039).

Finally, to test the impact of minor model misspecification, we fit an overparameterized ARD model to data simulated under equal rates. Under this model, performance was consistently lower than the ER model (Figure 2). The accuracy gap between joint and marginal methods widened, particularly at medium rates (0.5), where joint reconstruction maintained 30.0% accuracy compared to 26.5% for marginal reconstruction in two-state characters (McNemar’s χ² = 18.35, p < 0.0001). This difference was similarly pronounced for three-state characters at medium rates (McNemar’s χ² = 17.65, p < 0.0001). Even at low rates (0.25), the methods showed significant differences under the ARD model for both two-state (McNemar’s χ² = 11.26, p = 0.0008) and three-state characters (McNemar’s χ² = 10.26, p = 0.001). However, three-state character reconstruction maintained relatively high accuracy (joint: 63.9%, marginal: 61.7%). Thus, for both ER and ARD models, the joint ancestral state reconstruction performed equally or better than the marginal reconstruction.

### Incorporating uncertainty is crucial for capturing the true evolutionary history

To evaluate the performance of our uncertainty quantification approaches, we examined our method’s ability to capture true ancestral states. The uncertainty quantification methods showed substantial performance increases (Figure 2). By generating multiple plausible reconstructions proportional to their likelihood, our uncertainty intervals captured the true state configurations with high reliability, particularly under the ER model (97.8% for three-state, 97.0% for two-state at low rates). Even as point estimate accuracy declined with increasing rates (5.9% for the two-state model and 4.7% for the three-state model), uncertainty interval coverage remained relatively robust (31.5% for the two-state model and 36.3% for the three-state model at high rates under ER). When examining the impact of model choice, we found that misspecification had a moderate impact on uncertainty interval coverage, with ARD model coverage dropping to 25.4% at high rates for three-state characters.

Our results further demonstrate that joint uncertainty intervals significantly outperform marginal intervals in capturing true ancestral reconstructions. This performance difference is particularly pronounced at medium and fast rates, where there is inherently greater uncertainty in ancestral predictions. Furthermore, the increased accuracy occurs despite joint sampling requiring fewer ancestral samples than marginal sampling approaches. This is likely due to the sampling methodology differing between approaches. Joint intervals are constructed by sampling complete ancestral histories based on their likelihood, whereas marginal intervals are generated by independently sampling nodes according to their individual probabilities. For our simulations, sampling continued until either all unique histories had been exhausted or a maximum threshold of 1,000 reconstructions had been reached. Interestingly, only when uncertainty was relatively low did, we observe joint sampling requiring more alternative reconstructions than marginal sampling. This pattern reveals an important methodological insight. Cases with apparently low uncertainty, where researchers might have false confidence in a reconstruction, are precisely where joint intervals identify plausible alternatives that are difficult to detect through independent node sampling. Finally, one of the potentially most significant advantages of joint uncertainty intervals is their statistical properties. Joint intervals consistently center around the true ancestral states while providing narrower confidence bounds than marginal intervals. This suggests that joint sampling efficiently captures the most reasonable alternative reconstructions. Marginal sampling tends to include many improbable reconstructions that, on average, are not centered on the true ancestral history. These findings indicate that joint uncertainty quantification methods provide more accurate and efficient estimations of ancestral state uncertainty, particularly in scenarios where character evolution occurs at moderate to rapid rates.

### Reconstruction uncertainty reveals distinct patterns of resistance to last-resort antibiotics in *Klebsiella pneumoniae*

Alongside these simulations, we applied our methodology to study the evolution of antibiotic resistance. ASR has been extensively leveraged to identify genetic determinants and characterize the evolutionary trajectory of resistant lineages, largely due to detectable signatures of convergent evolution via spontaneous mutation or the acquisition of mobile genetic elements (11–13). Here, we applied our framework to characterize the evolution of resistance to last-line antibiotic, β-lactam/β-lactamase inhibitor (BL/BLI) combinations, in 412 carbapenem-resistant *Klebsiella pneumoniae* sequence type 258 isolates collected across a long-term acute care hospital network from 2014 to 2015 (27). Equipped with a preliminary understanding of the genetic basis (14), we evaluated whether reconstruction uncertainty would illuminate our understanding of the evolutionary dynamics of resistance to these promising antibiotics.

Joint reconstruction revealed the circulation of several resistant lineages, alongside a background of resistance acquired on terminal branches, suggestive of frequent *de novo* evolution of resistance (Figure 3A). The uncertainty quantification method revealed many alternative evolutionary histories of BL/BLI resistance, with over 900 alternative reconstructions identified (Figure 3B). As individually analyzing each of these possibilities is infeasible, we developed a clustering approach based on a novel distance metric for ancestral reconstructions (Materials and Methods). This clustering approach created a distance matrix based on the weighted differences between ancestral reconstructions by combining differences in the character state and their phylogenetic distribution. We then applied t-distributed stochastic neighbor embedding (t-SNE) (28) and a density-based clustering algorithm (29) to identify five distinct clusters of reconstruction states (termed ‘reconstruction axes’) for BL/BLI resistance (Figure 3B). We note that the exact number and composition of clusters can vary depending on the hyperparameters of the joint uncertainty analysis and t-SNE clustering implementation. However, our chosen methodology yielded the most consistent results among several tested alternatives. Nonetheless, each reconstruction axis captured distinct reconstructions that described unique emergence events, altering the distribution of trait transitions and continuation events across the phylogenetic tree (Table 1). That is not to say there are no general patterns consistent across all reconstructions. Indeed, a visualization of important nodes with substantial variation across reconstruction axes revealed substantial variation within specific clades (Figure 3AC). Reconstruction axes were inferred to have varying frequencies of resistance emergence, spread, and loss, predominantly due to variation in the ancestral node where resistance was acquired. Collectively, these findings support differential hypotheses regarding the evolutionary history of resistance, its associated fitness costs, and potential pathways for acquiring resistance to these antibiotics.

**Figure 3.**
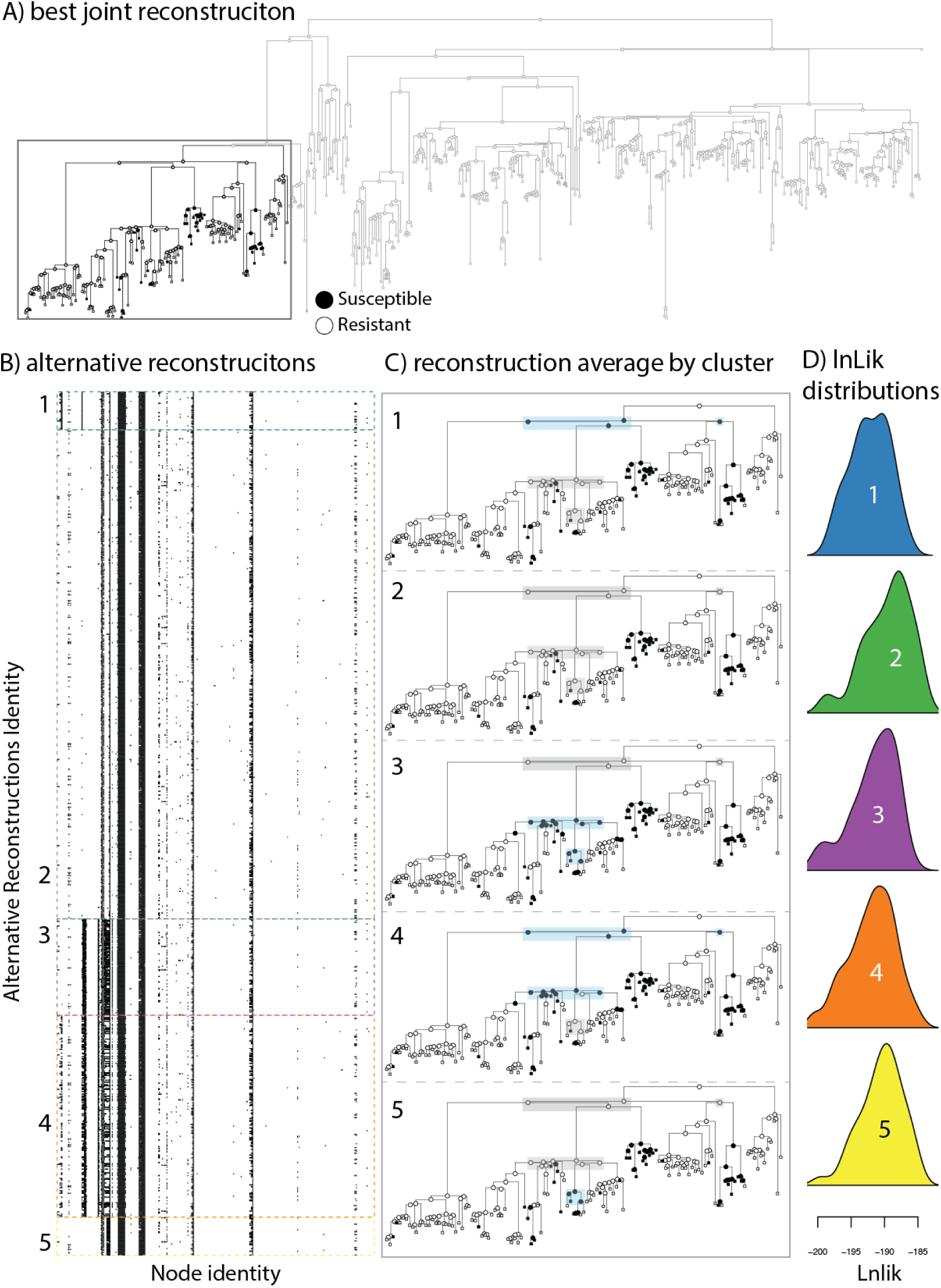
Reconstruction uncertainty illuminates differential hypotheses regarding the evolutionary history of antibiotic resistance. A) The maximum-likelihood phylogeny and best joint reconstruction of resistance to β-lactam/β-lactamase inhibitor among 412 carbapenem-resistant *Klebsiella pneumoniae* sequence type 258 isolates collected across a long-term acute care hospital network. Highlighted is the clade where most of the significant differences appear. B) Alternative reconstructions where node states are flattened. Each row represents an alternative reconstruction sequence, and each column corresponds to a different node index. Reconstructions have been grouped into various clusters found by our phylogenetically aware clustering algorithm. C) A representative reconstruction from each cluster, focusing on the most variable clade, as highlighted in panel A. Each reconstruction is chosen because it has the minimum average distance to all other reconstructions within the cluster. We have highlighted where key changes have occurred relative to the most likely joint reconstruction. A grey highlight indicates concordance with the most likely joint reconstruction and a blue highlight indicates where a susceptible reconstruction was estimated rather than resistance. D) The likelihood distribution of the reconstructions within each cluster.

**Table 1.**
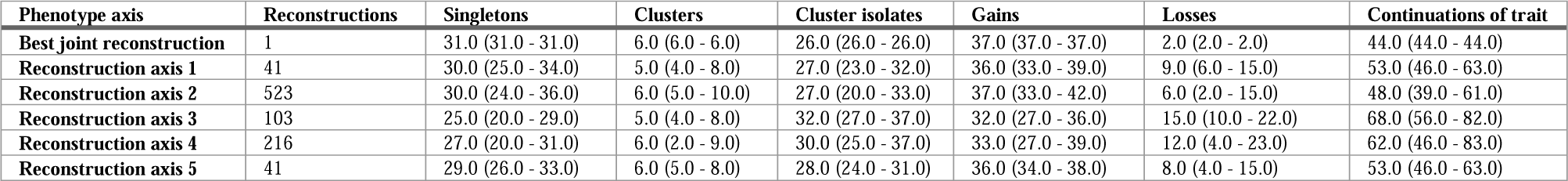
Summary statistics for phylogenetic reconstruction of antimicrobial resistance events across different reconstruction axis. Each row represents a different reconstruction axis, with the “Best joint reconstruction” showing the optimal joint reconstruction. Columns display: Singletons - number of phylogenetic singleton resistance events; Clusters - count of phylogenetic clusters of resistant isolates; Cluster isolates - number of isolates belonging to resistance clusters; Gain events - transitions from susceptible to resistant along edges of the phylogeny; Loss events - transitions from resistant to susceptible; Continuations - edges where the trait (resistance) persists. Values represent median estimates with ranges (minimum, maximum) across reconstruction iterations.

To test whether the different evolutionary histories found by our uncertainty framework influenced the appraisal of resistance-conferring genotypes, we used known resistance-associated genotypes as a form of truth testing. This analysis focused on several important genotypes implicated in BL/BLI resistance, including mutations to the *ompK36* porin, antibiotic efflux regulation (*ramR* efflux pump regulator and *ramA* efflux pump activator), target modification (penicillin-binding proteins [PBPs]), and antibiotic inactivation (*Klebsiella pneumoniae* carbapenemase [*bla_KPC_*] -containing plasmid, AA552, with elevated copy number).

We performed pairwise comparison of phenotype and genotype reconstructions, quantifying the number of events where the genotype and phenotype were gained on the phylogeny (Figure 4). As expected, whether genotypic data aligned more closely with resistance emergence events depended on which reconstructions were used (Figure 4A). Investigating the consistency of resistance acquisition with our genotypes revealed large differences in the consistency of synchronous transitions across reconstructions (Figure 4B), suggesting that select reconstructions may more favorably align with the dynamics of these resistance-associated genotypes. However, our pairwise integration of genotype and phenotype uncertainty also revealed several genotypes with minimal variation, with most reconstructions showing consistent, concordant predictions for trait evolution (Supplemental Materials). Finally, we investigated the utility of considering all genotypic reconstructions when explaining the evolution of resistance in this population. The proportion of phenotype gain events with at least one genotypic gain event varied across reconstruction axes. Notably, several reconstructions performed consistently better than the optimum joint reconstruction. Collectively, these findings suggest that evaluating uncertainty in phenotypic and genotypic reconstructions will improve our understanding of the evolutionary dynamics of a trait and will enable us to identify alternative hypotheses to consider for further testing.

**Figure 4.**
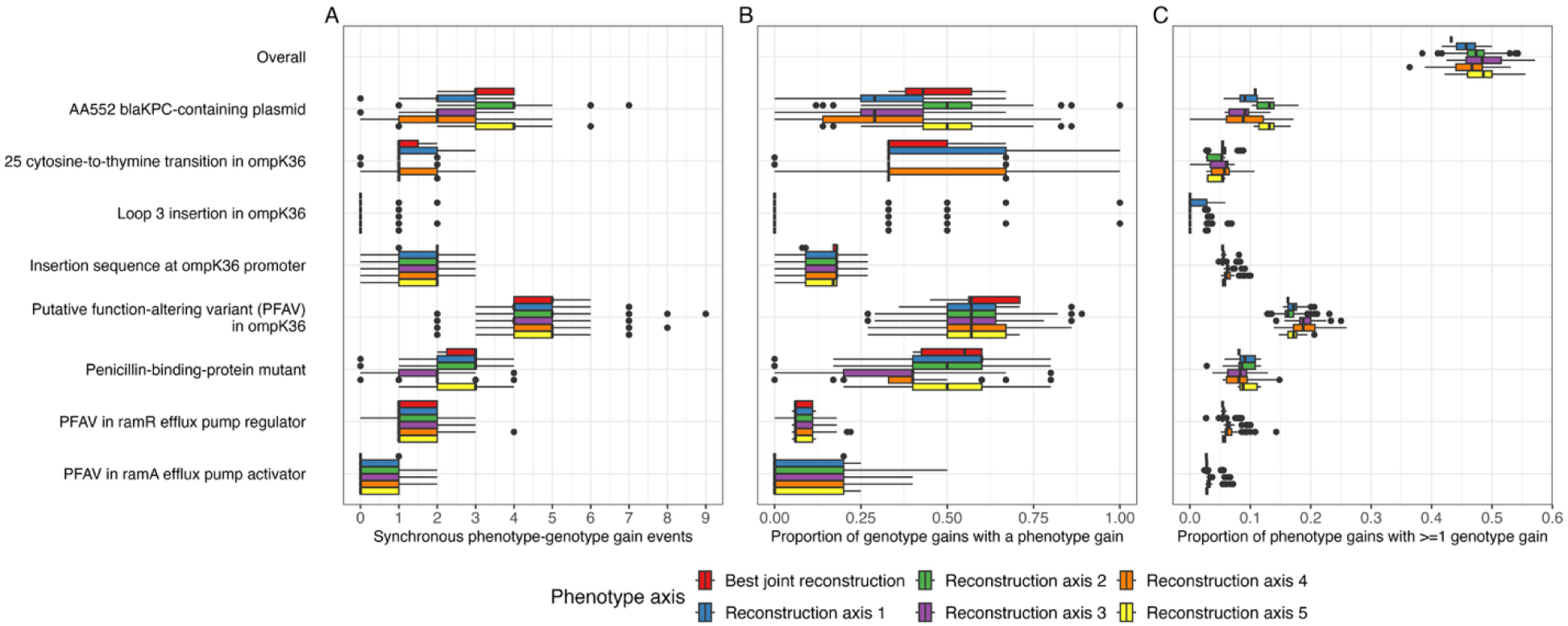
Comparison of synchronous phenotype-genotype transition events across different evolutionary reconstructions. We compared reconstructions of our phenotype, β-lactam/β-lactamase inhibitor combination resistance, and carbapenem resistance-associated genotypes to detect synchronous transitions across the phylogeny, evidence of genotype-phenotype correlations. The figure displays distributions of A) synchronous phenotype-genotype gain events B) proportion of genotype gain events corresponding with a phenotype gain event and C) the proportion of phenotype gain events in a reconstruction with >=1 genotype gain event, considered individually and collectively. The inferred strength of any single phenotype-genotype association depends on the reconstruction axis chosen.

## Discussion

### Reconstruction uncertainty impacts genotype-phenotype associations

Phylogenetic approaches to genome-wide association studies often use synchronous transition events as evidence for genotype-phenotype relationships (30, 31). Our analysis of BL/BLI resistance in *K. pneumoniae* demonstrates that relying on a single “best” reconstruction may significantly impact these inferences. By applying our uncertainty quantification approach, we identified multiple plausible reconstruction axes, each of which infers a distinct evolutionary history for antibiotic resistance. These alternative reconstructions had meaningful consequences for genotype-phenotype associations. Importantly, the frequency of synchronous phenotype-genotype-gain events differed substantially between reconstructions, with some axes placing genotype transitions downstream of phenotype emergence. While this initially appears to be a biologically implausible scenario (suggesting the phenotype preceded its genetic cause), such patterns may reflect more complex evolutionary dynamics. For instance, downstream genetic variation could represent fitness compensation, fine-tuning of resistance mechanisms, or resolution of fitness tradeoffs following the primary resistance-conferring mutation (32–34). Crucially, these alternative reconstructions enable domain experts to evaluate competing evolutionary scenarios using independent data, enhancing our understanding of resistance evolution.

Overall, these findings highlight a critical limitation of traditional approaches that select a single reconstruction. The “winner-take-all” strategy may lead researchers to overlook relevant genotypes simply due to the arbitrary selection of one evolutionary scenario among several equally plausible alternatives. For complex traits like antibiotic resistance, where multiple genetic mechanisms can produce similar phenotypes (35), this limitation is particularly problematic. By considering multiple plausible evolutionary scenarios when inferring associations, researchers can develop more robust models that better acknowledge the inherent uncertainty in ancestral state reconstruction.

An important avenue for future work is the development and application of correlated models that jointly infer the evolution of genotypes and phenotypes within a single framework. In our case, we modeled these components separately and then independently assessed relationships between events based on two distinct sets of ancestral reconstructions (one for genotypes and one for phenotypes). This approach requires considerable post hoc interpretation and multiplies uncertainty, since any inference depends on reconciling multiple plausible genotype reconstructions with multiple plausible phenotype reconstructions. By contrast, models of correlated evolution have been shown to outperform independent models and offer a direct way to assess evolutionary relationships between traits (36–40). For ancestral state reconstruction, such joint models would provide a single set of predictions, greatly reducing the interpretive complexity. Ultimately, this integrated approach would not only strengthen genotype–phenotype association tests but also yield a more coherent picture of how molecular and phenotypic traits evolve together.

### The case for joint ancestral state estimation with uncertainty

Ancestral state reconstruction can provide crucial insights into character evolution and serves as a foundation for many comparative analyses. However, it is also an unfortunate misnomer (22, 22, 23, 38, 41). The use of the word “reconstruction” implies more certainty than is often warranted, and some authors prefer the term ancestral state estimation instead (e.g., 41). Throughout this text, we have used the term “reconstruction” primarily for historical reasons, but ancestral states are literally estimates derived from predictions of our fitted model, making “estimation” the more accurate terminology. And when viewed as estimates, it should be clear that ancestral states will be uncertain because parameters cannot be perfectly known from a finite sample (25, 42, 43). This leads to the question: what kind of ancestral state uncertainty is most important for biological inferences?

In this text, we have made the case for joint reconstructions as the more biologically appropriate method for estimating ancestral states. This is because the unit of joint reconstructions is entire evolutionary histories rather than single nodes, and this better aligns with evolution as a highly contingent process (44). To our surprise, we found that considering the entire evolutionary history simultaneously via joint reconstruction also statistically outperformed marginal methods across all our simulation conditions. The field has traditionally relied on marginal reconstruction methods because they are a computationally efficient way to summarize all possible evolutionary histories (3, 45). The probabilities returned by marginal reconstructions do represent uncertainty, but only at a given node. The uncertainty provided does not directly represent the dependent relationships present in a phylogenetic tree (i.e., if an ancestor has a particular state, the descendant will not inherit it; all nodes are considered independently). In practice, marginal reconstructions mainly indicate where uncertainty is present, but they do not show how it is distributed across distinct evolutionary scenarios. A crucial limitation, therefore, is not just that they are less accurate, but that they fail to capture the covariation of states across the tree, which is often the basis of biological hypotheses.

This highlights the need not only to quantify uncertainty but also to do so in a way that preserves the structure of an entire evolutionary history. While joint reconstruction algorithms have existed for decades, they have traditionally focused on identifying only the single most likely history, leaving researchers with a point estimate that offers a false sense of certainty (24). What has been missing are methods to efficiently sample and summarize the full distribution of plausible joint histories. Our algorithms were developed to fill this gap. As our simulations demonstrate, the uncertainty captured by sampling joint histories is superior to that from marginal reconstructions. The resulting set of plausible histories is more likely to contain the true evolutionary path, and, perhaps more importantly, the distribution of these histories is centered around the true state configuration. In contrast, marginal sampling can produce many improbable combinations of states that, although possible at individual nodes, are globally unlikely, resulting in a less accurate representation of the true uncertainty.

Of course, using the full uncertainty of joint reconstructions presents its own set of practical challenges. If the output of an analysis is not one history but thousands, how can a researcher visualize, interpret, and draw conclusions from this information? The sheer number of alternative histories can be impossible to interpret directly. This is why we developed new analytical tools, such as phylogenetically aware clustering and dimensionality reduction techniques. These methods allow researchers to move beyond an overwhelming set of possibilities to a manageable and interpretable set of distinct “reconstruction axes” or representative histories. By identifying the major patterns of variation among all plausible scenarios, these approaches distill immense complexity into a small number of alternative, testable hypotheses about how a trait may have evolved. This makes the theoretical superiority of joint uncertainty empirically tractable.

Finally, it is essential to acknowledge that these methodological considerations are embedded within a broader debate about the fundamental reliability of ASR (46–50). Critics would argue that ancestral state estimation is little more than informed guesswork, as its claims are difficult to confirm or refute without direct evidence from fossils. While this critique is valid, we contend that they overlook that ancestral state estimates are a direct consequence of the evolutionary model. The model parameters are estimated from the data and have clear biological meaning (51). If we regard ASR as inherently unreliable, we are implicitly suggesting that the models of trait evolution on which they are based are similarly uninformative. In other words, to reject ASR as “useless” is not just to critique one application, but to undermine the broader modeling framework that underpins comparative methods in phylogenetics. A more productive approach is to acknowledge that ancestral reconstructions are model-based predictions subject to uncertainty, and to evaluate them with the same rigor we apply to any other inference derived from evolutionary models. To completely disregard these predictions because they are uncertain is to throw the baby out with the bathwater. By providing tools to explore competing evolutionary narratives, we move away from seeking a single “correct” reconstruction and toward a more robust framework for understanding the unobserved past.

## Conclusion

Our results demonstrate that joint ancestral state reconstructions, alongside methods to quantify and summarize uncertainty, provide more accurate and biologically meaningful insights than traditional marginal approaches. Unlike marginal methods, which focus on individual nodes in isolation, joint reconstructions treat entire evolutionary histories as the unit of inference. This perspective is more consistent with the contingent nature of evolution and better aligns with the kinds of questions biologists seek to answer. By summarizing several alternative histories as into major reconstruction axes, our framework allows uncertainty to be used to develop competing and testable hypotheses. Furthermore, our empirical analyses show how this approach reveals distinct evolutionary narratives with direct consequences for genotype–phenotype associations. Together, these advances establish a stronger foundation for ancestral state reconstruction and open the door to future models that integrate molecular and phenotypic evolution jointly.

## Materials and Methods

### Background for estimating the Probability of Ancestral States

The key distinction between a marginal and joint reconstruction is that the marginal provides probabilities for each node separately, while the joint considers the probability of complete configurations. In both cases, estimating these probabilities requires specifying a model of evolution along the tree. The most common choice is a continuous-time Markov model, which defines transition probabilities along branches (36). This model is fit to the discrete character data and phylogeny, and once optimized, is used for downstream ancestral state reconstructions. Joint reconstruction algorithms, as currently implemented (24), identify the most likely evolutionary scenario but do not provide any alternative hypotheses. In contrast, efficient marginal reconstruction algorithms provide uncertainty at each node (17) but give a more local view, which can be misleading if thinking about the full evolutionary history.

Consider an ancestral state reconstruction problem with *n* nodes and *k* possible states. Let *X_i_* represent the state of node *i*, where *i* = 1, ..., *n*. The state *X_i_* can take on any value from the set {1, ...*k*}. The marginal probability of node *i* being in state j is denoted as *P*(*X_i_* = *j*). This probability can be calculated by summing over all possible configurations of the other nodes:

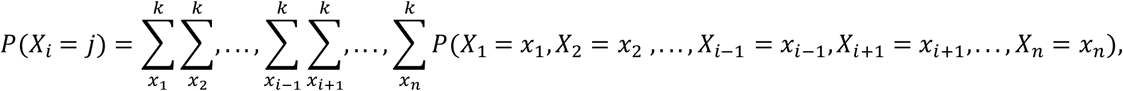

where each *x* ranges over the possible states {1, ..., *k*}. In other words, this equation says that to find the marginal probability of node *i* being in state *j*, we consider all possible combinations of states for the other nodes and sum the probabilities of each of those complete configurations. In contrast, the joint probability considers the probability of a specific configuration of states across all nodes. Let *x* = *x*_1_, ..., *x_n_* represent a configuration specifying the state of each node. The joint probability of a configuration *x* is denoted *P* (*X* = *x*) = *P*(*X*_1_ = *x*_1_, *X*_2_, ..., *X_n_* = *x_n_*).

### Algorithms for uncertainty quantification

#### Algorithm I: Stochastic sampling

Following Pupko et al. (2000), this algorithm proceeds by visiting nodes in a postorder fashion. Two sets of values are being calculated during this postorder traversal. First, a matrix of likelihoods, *L*, which specify the best reconstruction of a particular subtree conditioned that the parent of the subtree has a particular state *i*. Second, a matrix of character states, *C*, assigned to a particular node under the conditions of the optimal subtree. These two matrices have the same dimensions with the number of rows equal to the sum of the number of nodes and tips, and columns equal to the number of states. Tips are initialized with a likelihood of *e^Qt^*, where *t* is the branch length between the ancestor and the focal tip and *Q* is the instantaneous rate matrix. The most likely state is simply the observed state. After likelihoods for all tips are calculated, the next step is to visit nonroot internal nodes that have had both descendents visited. Under this algorithm, the likelihood that node *x* is in state *i* is *L_x_*(*i*) = *max_j_*(*P_ij_*(*t_x_*) × *L_y_*(*j*) × *L_z_*(*j*)) where *t_x_* is the branch length from the parent of node *x* to node *x*, and *L_y_*(*j*) is the likelihood of the best reconstruction of the subtree specified by node y given the parent node is assigned state *i*. *C_x_*(*i*) is then the value of j which corresponds to the above maximum. However, it is possible to modify this algorithm such that we sample states proportional to *P_ij_* (*t_x_*) × *L_y_*(*j*) × *L_z_*(*j*) instead of taking the maximum. Following this procedure would no longer result in calculating the the best reconstruction, but rather a sample from the possible subtree reconstructions when the parent has a particular state *i*. Because the joint probabilities represent a global reconstruction, each sample that differs from the maximum will alter the overall reconstruction and likelihoods. Sampling in proportion to *P_ij_* (*t_x_*) × *L_y_*(*j*) × *L_z_*(*j*) ensures that only plausible reconstructions are being considered and that alternative reconstructions are explored proportional to their overall likelihood.

The algorithm concludes in the same way regardless of whether one is interested in the most likely joint reconstruction or a sample. Once all non-root nodes have been visited the root is calculated in much the same way except that the combined descendent likelihoods at the root node are multiplied by the root prior. The final step is to proceed towards the tips reconstructing node *x* by choosing *C_x_*(*i*) where *i* is the most likely state of the ancestor. Also note that the formulas being presented are explicitly for bifurcating trees, but the multiplication can be extended to account for polytomies.

#### Algorithm II: Exact computation

Our alternative implementation proceeds by first calculating the marginal ancestral states along with the conditional probabilities at each node for each state. Then, the optimal joint likelihood configuration is calculated, along with its corresponding likelihood score. Because this has the highest probability among all configurations, it will be used to compare alternative configurations. Then, we conduct a single postorder traversal (from the tips to the root) through the tree. Our goal is to add to the best joint configuration, the set of other plausible reconstructions. Because the total number of alternative reconstructions scales by *n_nodes_^n_states_^*, we cannot calculate every alternative but for very small trees and state spaces. To record plausible reconstructions, we aim to efficiently filter out unreasonable alternatives, in terms of the distance of the likelihood from the best configuration, as we move from the tips to the roots. At each node, as we proceed, we build the reasonable alternatives for the subtending edges to that node such that once we have reached the root, we have the full set of alternatives. For each tip, we begin a plausible joint configuration that includes the observed states, and the corresponding likelihood. As we proceed to the parent node, for each state with a lnL for the marginal probability that is less than 2 away from the best lnL of the states at that node, we calculate the likelihoods of these states to each configuration of each child to this node. If the corresponding likelihood is no greater than 2 lnL worse than the best configuration, we add this state and likelihood to the child configuration. This proceeds until we come to the root where final likelihoods have composition incorporated.

### Analyzing patterns in alternative reconstructions

Both uncertainty algorithms produce multiple potential state reconstructions. It is infeasible to consider all the possible alternatives as the number of possible ancestral state reconstructions will scale exponentially with tree size and state space. While individual visualization of reconstructions is feasible when uncertainty is low or not considered, scenarios with substantial uncertainty will require an alternative analysis scheme. To address this, we developed a clustering-based approach to identify distinct patterns among a sample of *M* alternative joint reconstructions while preserving biologically relevant variation. The method employs a three-stage analysis pipeline: distance calculation, dimensionality reduction, and density-based clustering.

First, we quantified the dissimilarity between all M alternative ASRs. Each reconstruction, *R_k_*(*for k* = 1, ..., *M*), is a vector of character states (*S*_*k*,1_, *S*_*k*,2_, ..., *S_k, N_internal__* assigned to the *N_internal_* internal nodes of the phylogeny. Pairwise distances, *d*(*R_i_, R_j_*), were computed using a custom phylogenetically informed metric (implemented as phylo_aware_dist_local in corHMM). For any two reconstructions *R_i_* and *R_j_*, let N_*diff*_ = {*n*|*S_i,n_* ≠ *S_j,n_*} be the set of internal nodes where their assigned states differ. If the number of differing nodes, |N_*diff*_|, is less than or equal to one, *d*(*R_i_*, *R_j_*) = 0. Otherwise, the distance is defined as:

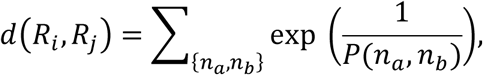

where the sum is over all unique pairs of distinct nodes {*n_a_, n_b_*} within N_*diff*_, and *P* (*n_a_, n_b_*) is the patristic distance between nodes *n_a_* and *n_b_* on the phylogeny (calculated using ape::dist.nodes on a tree with all branch lengths set to 1). This metric emphasizes sets of co-occurring state differences at phylogenetically proximate nodes, assigning higher dissimilarity scores when multiplied differences are concentrated in specific areas of the phylogeny. This will yield *D*, an *M* × *M* symmetric distance matrix.

Second, the distance matrix, *D*, is used as input for t-distributed Stochastic Neighbor Embedding (t-SNE) to reduce dimensionality (28, 52). Default t-SNE parameters were used (via Rtsne::Rtsne), except for enabling custom distance matrix input (is_distance = TRUE). This transformation projects the high-dimensional space of reconstructions onto a two-dimensional manifold, attempting to preserve the local structure (i.e., similarities between reconstructions). Finally, Density-Based Spatial Clustering of Applications with Noise (DBSCAN; 29) was applied to the two-dimensional t-SNE coordinates to identify clusters of similar ASRs. The DBSCAN parameters were determined following established guidelines (29). The minimum number of points required to form a dense region (minPts) was set to 15, representing roughly 1.5% of the total number of reconstructions. The neighborhood radius (eps) was subsequently determined by examining a k-nearest neighbor distance plot (dbscan::kNNdistplot) for *k* = *minPts* - 1. The value of eps was chosen at the “knee” of this plot, which indicates a suitable threshold to distinguish informative points from noise.

To identify diagnostic nodes distinguishing between clusters, we developed a node importance metric for each node and each cluster (implemented as node_importance_by_cluster in corHMM): *I* = (*P_w_*)^2^ * (1 - *P_b_*)^2^, where *I* represents node importance, *P_w_* is the proportion of matching ancestral states within a cluster, and *P_b_* is the proportion of matches between clusters, calculated using Hamming distances from the modal within-cluster reconstruction. Nodes exceeding an importance threshold of 0.5 were considered diagnostic for their clusters. The plot_alternative_jnts function in corHMM visualizes these alternative reconstructions by displaying the proportion of reconstructions with each state at each node within a cluster (53). While these proportional displays resemble marginal reconstruction pie charts, their interpretation differs. The clustering process has already identified and separated the major patterns of variation across the tree (global differences). Therefore, the pie charts within each cluster show only the remaining variation at individual nodes (local differences) among similar reconstruction scenarios. This separation of global and local patterns allows us to examine node-specific variation within each distinct reconstruction pattern, rather than averaging across all possible scenarios and potentially conflating patterns from distinct global scenarios (as in the marginal).

### Simulation protocol for ASR evaluation

We conducted a series of simulations to assess the performance of joint and marginal ancestral state reconstruction (ASR) methods. Trees were simulated using the geiger R package (54) with a fixed number of taxa (ntaxa=101 and nnode=100) and a tree height of 1. Character data were simulated using the corHMM R package (53) under a continuous-time markov model with equal transition rates between states (ER model). To investigate the impact of transition rate and number of states on ASR performance, we varied the rate parameter (0.25, 0.5, 1.0) and the number of states (k=2,3). For each combination of parameters, we simulated 1000 replicate datasets. This design allows us to evaluate the methods across a range of scenarios known to influence the performance of ASR methodology (38, 55, 56).

For each simulated dataset, we fit two models using corHMM: an equal-rates (ER) model, matching the generating model, and an all-rates-different (ARD) model. While the ARD model is known to perform poorly for ASR tasks (23), it is commonly used in practice and therefore we include it for completeness. Ancestral states were reconstructed using the ancRECON function with rate parameters estimated from the data. Additionally, we calculated the exact likelihood of each ancestral state configuration to determine equivalence between scenarios. Because marginal reconstructions provide uncertainty at each node, we take the maximum probability at each node to construct the most likely scenario under a marginal ASR.

To compare the performance of joint and marginal ASR, we first determine the true ancestral state configuration for each simulated dataset. We then evaluated whether the joint and marginal reconstructions correctly identified this configuration. We summarized the results using a confusion matrix reporting the proportion of simulations in which: (1) both joint and marginal methods were correct, (2) both were incorrect, (3) marginal was correct and joint was incorrect, and (4) joint was correct and marginal was incorrect for each of our simulation scenarios. Finally, to assess the reliability of our uncertainty quantification, we calculated the proportion of the time that the true ancestral states were captured by our uncertainty intervals. Additionally, we calculate a likelihood distribution of possible joint reconstructions based on our uncertainty quantification methods and examine how often the exact likelihood of the true ancestral states is found within this distribution.

### Ancestral state analysis of BL/BLI Resistance in *K. pneumoniae*

We applied our uncertainty methodology to evaluate antibiotic resistance patterns in *Klebsiella pneumoniae*. This analysis focused on resistance to recently introduced β-lactam/β-lactamase inhibitor (BL/BLI) antibiotics within a densely sampled population of 412 carbapenem-resistant *K. pneumoniae* isolates belonging to the epidemic lineage, sequence type 258 (27). Briefly, carbapenem-resistant *K. pneumoniae* identified from clinical isolates among long-term acute care patients were subjected to short-read whole-genome sequencing on an Illumina HiSeq 2500 instrument. A Gubbins-recombination filtered maximum-likelihood phylogeny was reconstructed using IQ-TREE version 2.4.0 using the GTR+F+ASC+R2 substitution model. IQ-TREE was performed using default parameters, other than the requirement of 500 unsuccessful iterations to halt reconstruction. A total of 5,000 bootstraps were performed. More information on the sample collection can be found elsewhere (27).

Phenotypic resistance was measured for two BL/BLI agents, meropenem-vaborbactam and imipenem-relebactam, using the SENSITITRE broth microdilution (Thermo Fisher; 55). Resistance to BL/BLI agents was determined as a composite of resistance to these agents, per 2021 Clinical and Laboratory Standard Institute interpretative criteria.

The evolutionary history of BL/BLI resistance was evaluated using ancestral state reconstruction. First, joint reconstruction was performed using corHMM under an equal rates model, with root probability estimated using the “maddfitz” procedure. Next, uncertainty in joint reconstruction was evaluated using corHMM’s *compute_j*oint_unc() with a batch size of 100 and a max sampling of 1000.

To illustrate the influence of ancestral state reconstructions classifications on evolutionary inferences, we leveraged the open-source R package, phyloAMR (15; https://github.com/kylegontjes/phyloAMR). To trace the ancestral state of each isolate, we applied a tree traversal algorithm, asr_cluster_detection(). Using ancestral states, the approach walks from the root to every tip, looking for evidence of trait continuation (e.g., susceptible to susceptible), gain (e.g., susceptible to resistance), or loss (e.g., resistance to susceptible). Next, the algorithm takes each trait-containing tip and walks upward on the tree to classify strains as either phylogenetic singletons or members of a phylogenetic cluster associated with the trait. Trait-containing tips with gain events inferred at the tip were classified as phylogenetic singletons if there was no history of ancestral resistance deeper in the tree. However, tips with a trait and a gain event at the tip and a trait reversion event at their parent node were eligible for classification as a cluster isolate. Tips were classified as members of a phylogenetic cluster if their first inferred ancestral gain event was shared with at least one additional isolate with a trait. Isolates with the trait that did not share a gain event with any other resistant isolate were classified as phylogenetic singletons. Lineages, where all isolates belonged to one patient, were not considered evidence of phylogenetic clustering and were classified as redundant singletons. The algorithm’s two identified states, singletons and clusters, permit distinguishing between *de novo* emergence of a trait and the transmission of trait-containing lineages, respectively. This algorithm was leveraged to test the inferences of resistance dynamics, notably number of phylogenetic events, gain/loss events, and size of clusters, across the alternative reconstruction axes inference identified using the clusters from the uncertainty algorithm.

To contextualize antibacterial resistance, we leveraged a collection of known carbapenem resistance-associated genotypes and a dominant *bla*_KPC_-containing plasmid previously described elsewhere (14). Individually identified single-nucleotide polymorphisms, insertion/deletions, and insertion sequences were simplified through binary classification (e.g., present/absent). Specifically, genetic variants were grouped by gene to enhance detection power for genotype-phenotype associations. For example, multiple non-synonymous mutations in the *ompK36* porin were consolidated into a single binary trait, as individual mutations might be too rare for meaningful statistical analysis.

Finally, we investigated the temporal relationship between genotype and phenotype transitions using the joint uncertainty and tree traversal algorithms. Phylogenetic transitions in genotype and phenotype are often used in genome-wide association studies to identify genotype-phenotype correlations (58, 59). We hypothesized that ancestral state differences influence the robustness of identified genotype-phenotype associations. To test this hypothesis, we evaluated the frequency of synchronous gain events of BL/BLI resistance and our resistance-associated genotypes using <u>phyloAMR’s synchronous_transition(</u>) algorithm. We also evaluated the consistency of genotype gain events with resistance acquisition and the proportion of phenotypic gain events where at least one genotype reconstruction inferred a genotype gain event, individually and collectively across all genotypes. All code for this empirical case study is hosted at https://github.com/kylegontjes/joint-uncertainty-ms.

## Acknowledgments

E.S., and K.J.G. were supported by National Institute of Allergy and Infectious Diseases (NIAID) grant 5R01AI148259. J.B., K.J.G., S.A.S., and E.S. were supported by the NIAID U19AI181767. S.A.S. was also supported by National Science Foundation (NSF) IntBio 2217116. K.J.G. reports training fellowships from the National Human Genome Research Institute (T32-HG000040) and NIAID (F31-AI186288).

## Supplementary Figures

**Figure S1.**
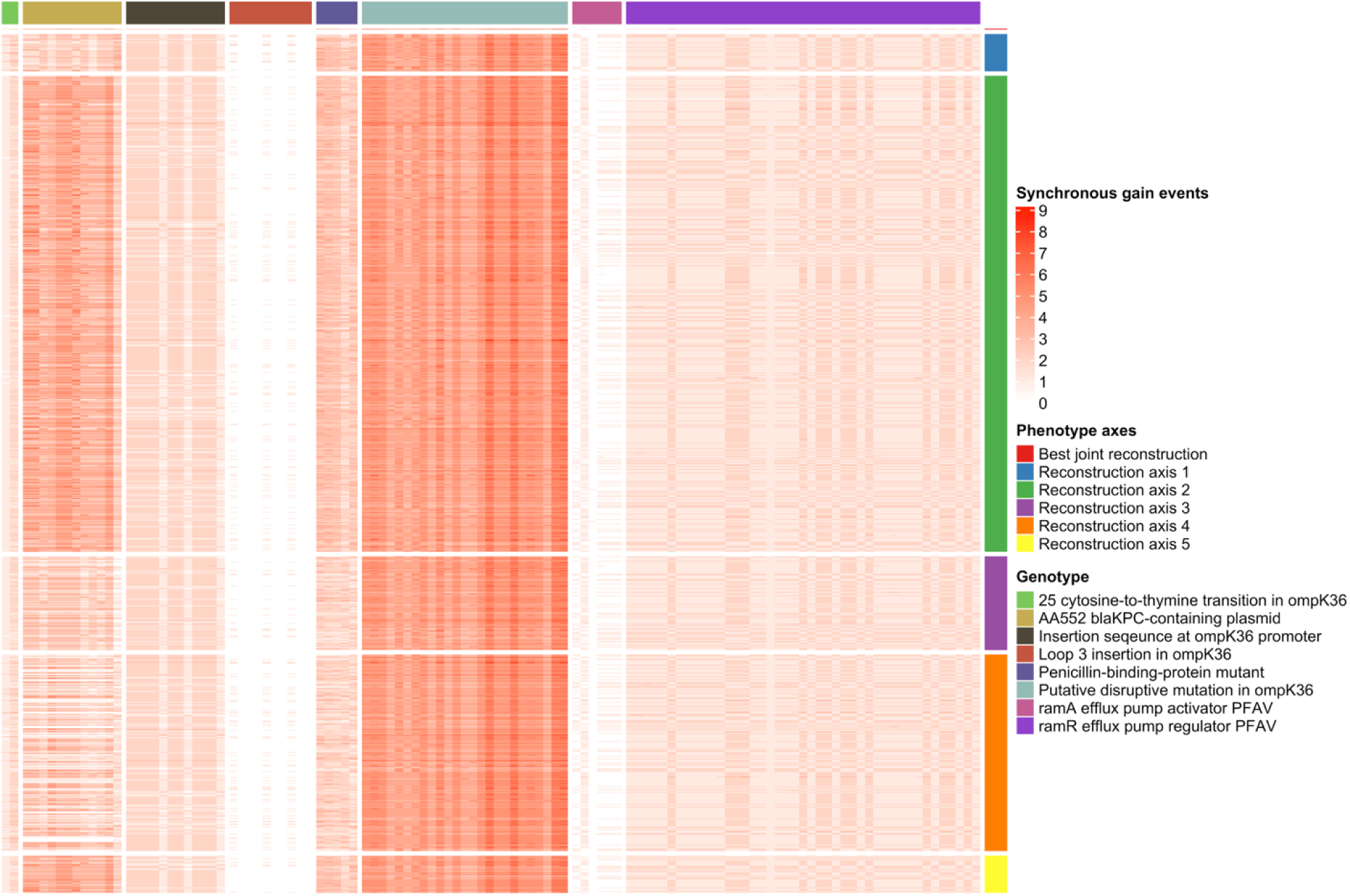
Heatmap of concordant phenotype-genotype gain events across all pairwise model reconstructions. Each cell represents a single pairing of a phenotype reconstruction (row) and a genotype reconstruction (column). The intensity of the color corresponds to the number of concordant transitions. Rows are grouped by the major phenotype axis to which the reconstruction belongs (annotation on right), and columns are grouped by the resistance genotype (annotation on top). This visualization highlights how the strength of association, measured by synchronous events, varies across the landscape of model uncertainty.

**Figure S2.**
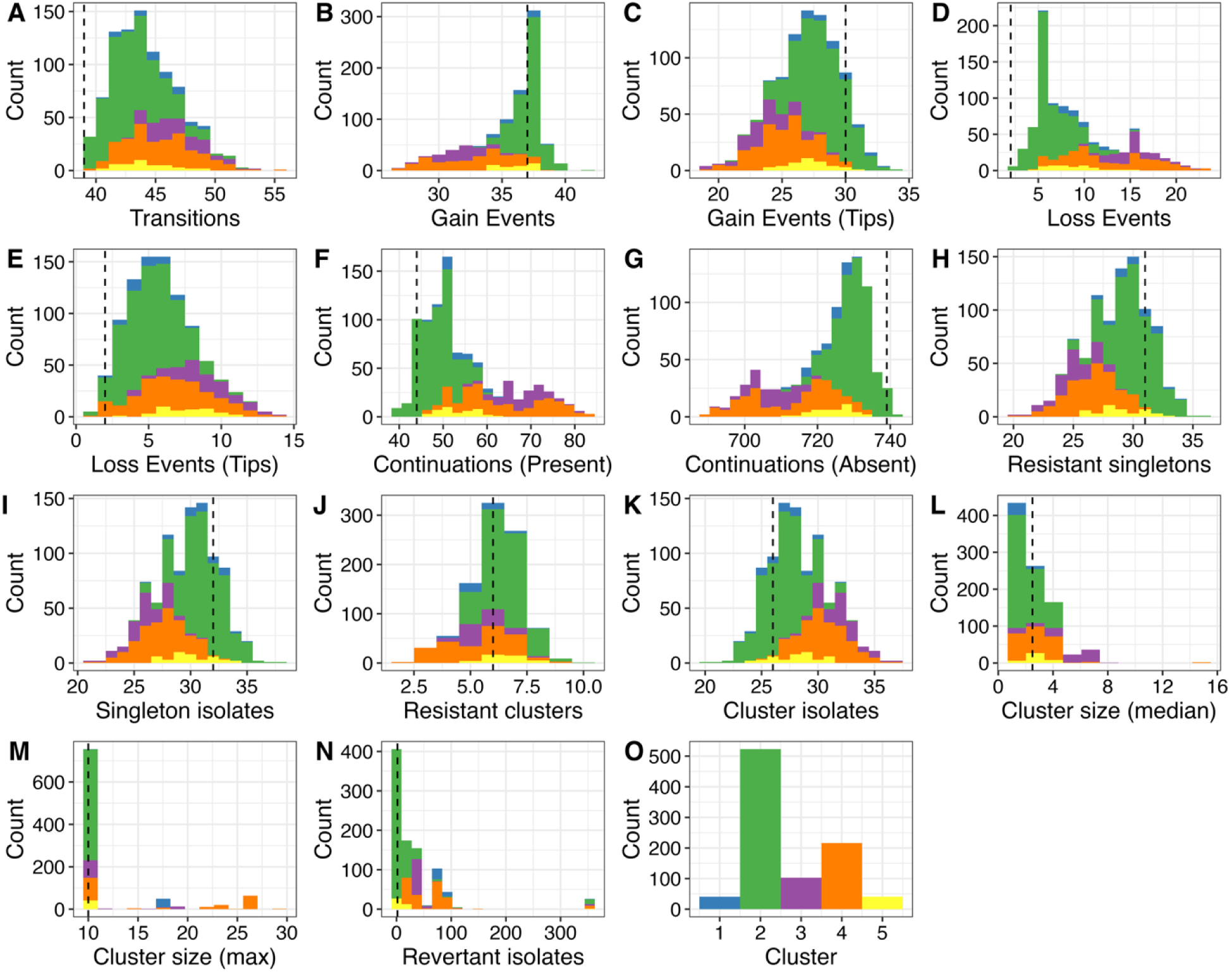
Distribution of evolutionary metrics across different ancestral reconstruction axes for resistance to β-lactam/β-lactamase inhibitor combinations. Each panel shows a histogram of a specific evolutionary metric, such as the number of gain events (B, C) or loss events (D, E), trait continuation and loss events (F,G), alongside the frequency and size of phylogenetic emergence events (H-M) and loss events (N) derived from the full set of alternative reconstructions. Colors correspond to the major reconstruction axis (cluster) to which each reconstruction belongs. Dashed vertical lines indicate the value for the best joint reconstruction model.

**Figure S3.**
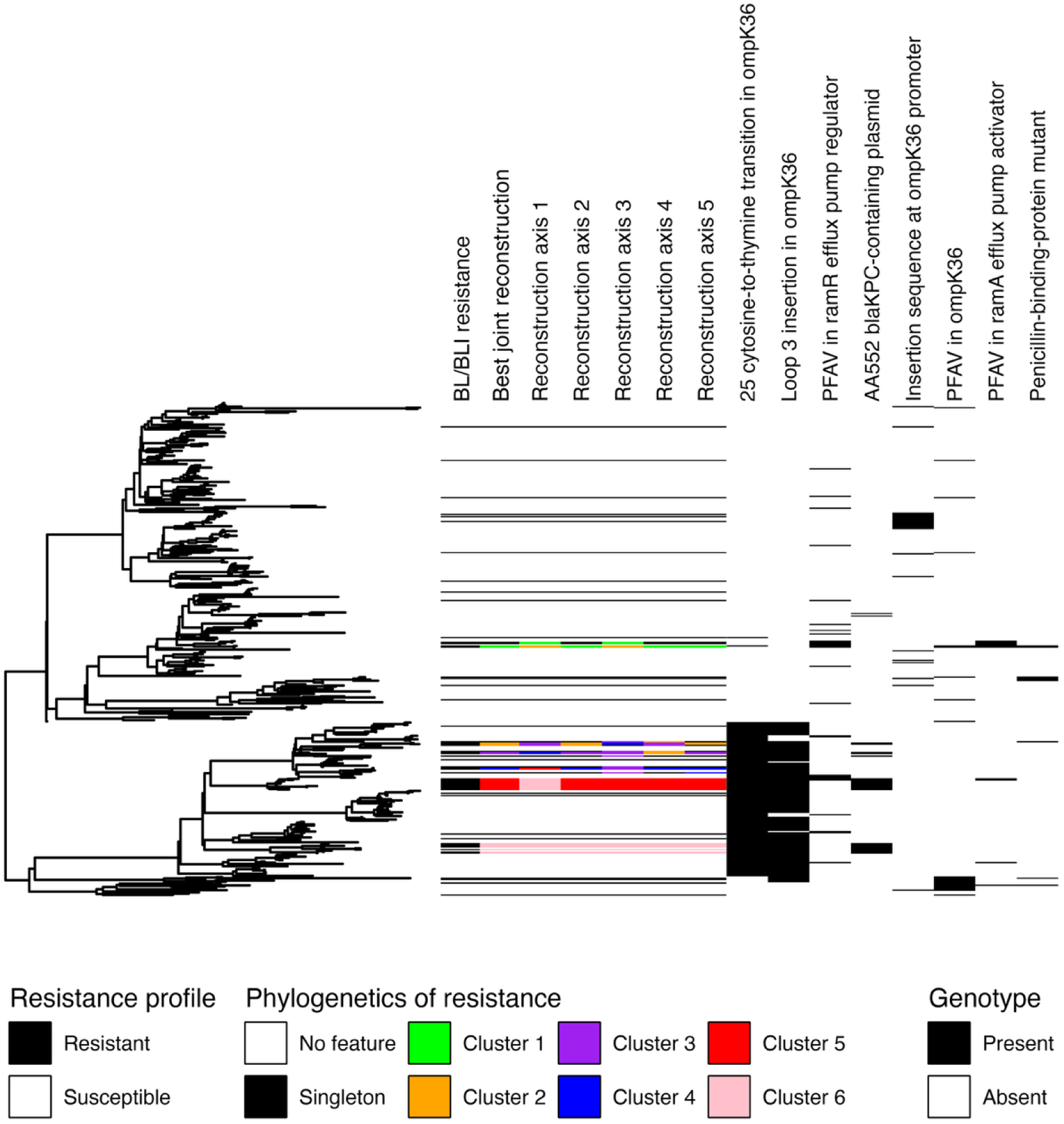
Phylogenetic distribution of resistance phenotypes, genotypes, and inferred evolutionary dynamics. The central phylogeny is annotated with multiple data panels. From left to right: (1) The measured resistance profile (Resistant/Susceptible) for each isolate. (2) The phylogenetics of β-lactam/β-lactamase inhibitor resistance as inferred by the best joint reconstruction model and five alternative reconstruction axes, showing whether resistance at each tip is classified as a “singleton” (i.e., independent emergence) or part of a phylogenetic “cluster” (i.e., circulation of resistant lineage). (3) A heatmap showing the presence/absence of key carbapenem resistance-associated genotypes. This visualization highlights how different reconstruction axes can lead to different interpretations of resistance emergence, as seen in the varying patterns of singletons and clusters for the same set of resistant isolates.

**Figure S4.**
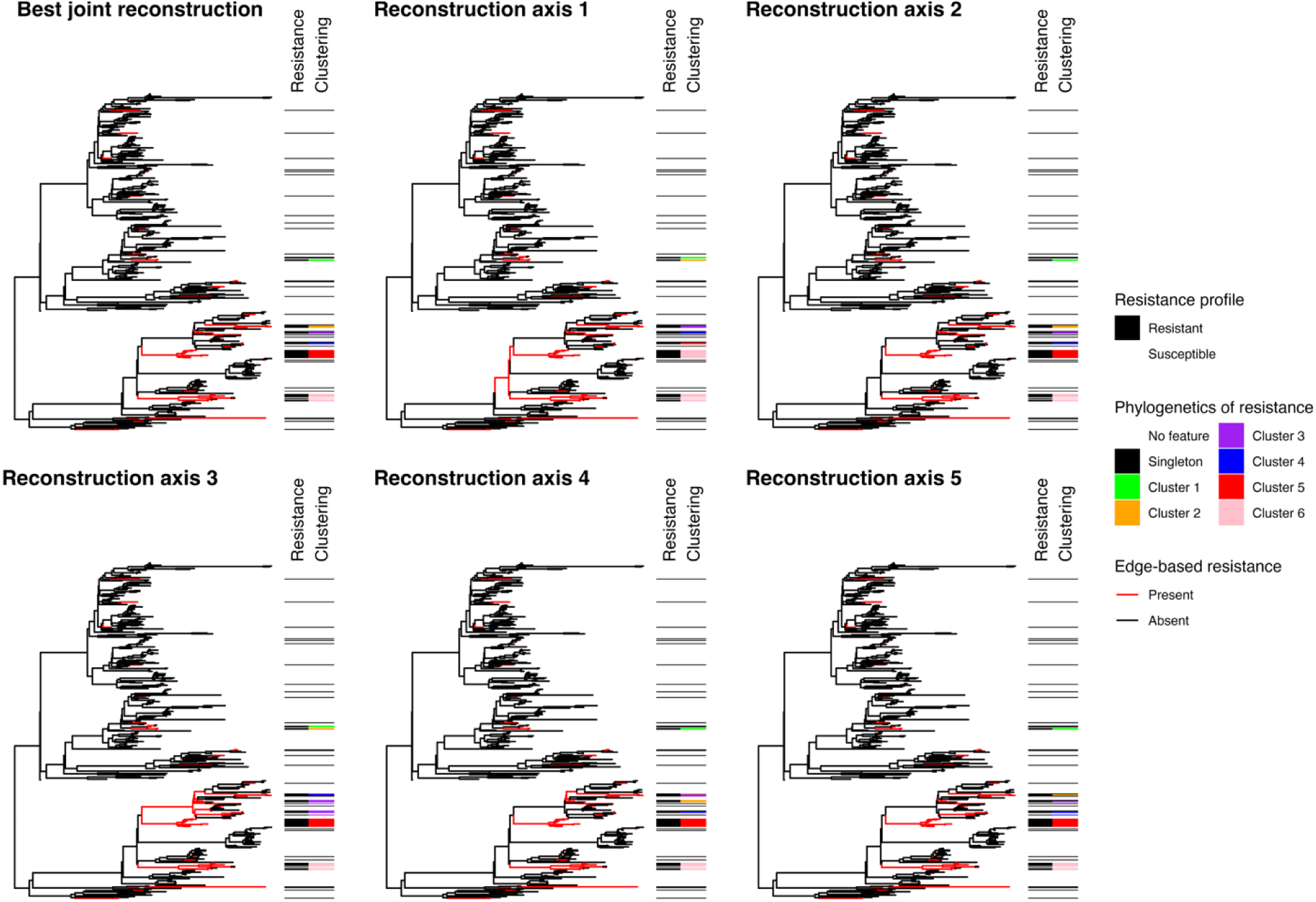
Inferred evolutionary histories of β-lactam/β-lactamase inhibitor resistance for six representative reconstruction models. Each panel shows the phylogeny with branches colored by the inferred presence (red) or absence (black) of resistance for a specific model. For each alternative reconstruction axis, the reconstruction with the lowest log-likelihood value was utilized. The annotations show the observed resistance profile and the resulting classification of resistant isolates into singletons or transmission clusters. The variation across panels highlights the uncertainty in identifying specific gain events and transmission dynamics.

## Supplementary Tables

**Table S1.** Summary statistics for evolutionary dynamics of resistance to β-lactam/β-lactamase inhibitor combinations across reconstruction axes. The table reports the median and range (minimum-maximum) for clustering and transition metrics for each of the six representative reconstruction axes.

| Phenotype axis | Reconstructi<br>ons | Phylogenetic<br>events | Singletons | Singleton<br>isolates | Clusters | Cluster<br>isolates | Cluster size<br>(median) | Cluster size<br>(max) | Transitions | Gain<br>events | Gains (tip) | Loss<br>events | Losses<br>(tip) | Continuations<br>of trait |
| --- | --- | --- | --- | --- | --- | --- | --- | --- | --- | --- | --- | --- | --- | --- |
| Reconstruction<br>axis 1 | 41 | 35.0 (33.0 -<br>38.0) | 30.0 (25.0 -<br>34.0) | 31.0 (26.0 -<br>35.0) | 5.0 (4.0 -<br>8.0) | 27.0 (23.0 -<br>32.0) | 2.0 (2.0 - 3.0) | 17.0 (10.0 -<br>17.0) | 45.0 (41.0 -<br>49.0) | 36.0 (33.0 -<br>39.0) | 28.0 (24.0 -<br>33.0) | 9.0 (6.0 -<br>15.0) | 5.0 (2.0 -<br>8.0) | 53.0 (46.0 -<br>63.0) |
| Reconstruction<br>axis 2 | 523 | 37.0 (33.0 -<br>41.0) | 30.0 (24.0 -<br>36.0) | 31.0 (25.0 -<br>38.0) | 6.0 (5.0 -<br>10.0) | 27.0 (20.0 -<br>33.0) | 2.0 (2.0 - 4.0) | 10.0 (10.0 -<br>10.0) | 43.0 (39.0 -<br>52.0) | 37.0 (33.0 -<br>42.0) | 28.0 (21.0 -<br>34.0) | 6.0 (2.0 -<br>15.0) | 5.0 (1.0 -<br>12.0) | 48.0 (39.0 -<br>61.0) |
| Reconstruction<br>axis 3 | 103 | 31.0 (27.0 -<br>34.0) | 25.0 (20.0 -<br>29.0) | 26.0 (21.0 -<br>31.0) | 5.0 (4.0 -<br>8.0) | 32.0 (27.0 -<br>37.0) | 5.0 (2.0 - 8.5) | 10.0 (10.0 -<br>19.0) | 46.0 (43.0 -<br>53.0) | 32.0 (27.0 -<br>36.0) | 24.0 (19.0 -<br>29.0) | 15.0 (10.0 -<br>22.0) | 9.0 (4.0 -<br>14.0) | 68.0 (56.0 -<br>82.0) |
| Reconstruction<br>axis 4 | 216 | 32.0 (26.0 -<br>37.0) | 27.0 (20.0 -<br>31.0) | 28.0 (21.0 -<br>33.0) | 6.0 (2.0 -<br>9.0) | 30.0 (25.0 -<br>37.0) | 3.0 (2.0 - 15.5) | 15.0 (10.0 -<br>29.0) | 45.0 (40.0 -<br>55.0) | 33.0 (27.0 -<br>39.0) | 25.0 (19.0 -<br>30.0) | 12.0 (4.0 -<br>23.0) | 6.0 (1.0 -<br>14.0) | 62.0 (46.0 -<br>83.0) |
| Reconstruction<br>axis 5 | 41 | 35.0 (33.0 -<br>38.0) | 29.0 (26.0 -<br>33.0) | 30.0 (27.0 -<br>34.0) | 6.0 (5.0 -<br>8.0) | 28.0 (24.0 -<br>31.0) | 3.0 (2.0 - 4.0) | 10.0 (10.0 -<br>10.0) | 44.0 (41.0 -<br>49.0) | 36.0 (34.0 -<br>38.0) | 27.0 (24.0 -<br>31.0) | 8.0 (4.0 -<br>15.0) | 8.0 (3.0 -<br>11.0) | 53.0 (46.0 -<br>63.0) |
| Best joint<br>reconstruction | 1 | 37.0 (37.0 -<br>37.0) | 31.0 (31.0 -<br>31.0) | 32.0 (32.0 -<br>32.0) | 6.0 (6.0 -<br>6.0) | 26.0 (26.0 -<br>26.0) | 2.5 (2.5 - 2.5) | 10.0 (10.0 -<br>10.0) | 39.0 (39.0 -<br>39.0) | 37.0 (37.0 -<br>37.0) | 30.0 (30.0 -<br>30.0) | 2.0 (2.0 -<br>2.0) | 2.0 (2.0 -<br>2.0) | 44.0 (44.0 -<br>44.0) |

**Table S2.** Summary statistics for phenotype-genotype transitions. Median and range (minimum-maximum) of concordant and discordant transition counts for all tested genotypes across each of the six representative reconstruction axes.

| Phenotype axis | AA552 *bla*<sub>KPC</sub>-containing plasmid synchronous gains | 25 cytosine-to-thymine transition in *ompK36* synchronous gains | Loop 3 insertion in *ompK36* synchronous gains | Insertion sequence at *ompK36* promoter synchronous gains | Putative disruptive mutation in *ompK36* synchronous gains | Penicillin-binding-protein mutant synchronous gains | *ramR* efflux pump regulator PFAV synchronous gains | *ramA* efflux pump activator PFAV synchronous gains |
| --- | --- | --- | --- | --- | --- | --- | --- | --- |
| Reconstruction axis 1 | 2.0 (0.0 - 4.0) | 1.0 (1.0 - 3.0) | 0.0 (0.0 - 2.0) | 2.0 (0.0 - 3.0) | 4.0 (3.0 - 7.0) | 3.0 (0.0 - 4.0) | 1.0 (1.0 - 2.0) | 0.0 (0.0 - 1.0) |
| Reconstruction axis 2 | 4.0 (1.0 - 7.0) | 1.0 (0.0 - 2.0) | 0.0 (0.0 - 1.0) | 2.0 (0.0 - 3.0) | 5.0 (2.0 - 9.0) | 3.0 (0.0 - 4.0) | 1.0 (0.0 - 3.0) | 0.0 (0.0 - 2.0) |
| Reconstruction axis 3 | 2.0 (0.0 - 4.0) | 1.0 (0.0 - 2.0) | 0.0 (0.0 - 1.0) | 2.0 (0.0 - 3.0) | 4.0 (2.0 - 7.0) | 2.0 (0.0 - 4.0) | 1.0 (1.0 - 3.0) | 0.0 (0.0 - 2.0) |
| Reconstruction axis 4 | 2.0 (0.0 - 5.0) | 1.0 (0.0 - 3.0) | 0.0 (0.0 - 2.0) | 2.0 (0.0 - 3.0) | 5.0 (2.0 - 8.0) | 2.0 (0.0 - 4.0) | 1.0 (1.0 - 4.0) | 0.0 (0.0 - 2.0) |
| Reconstruction axis 5 | 4.0 (1.0 - 6.0) | 1.0 (1.0 - 2.0) | 0.0 (0.0 - 1.0) | 2.0 (0.0 - 2.0) | 5.0 (2.0 - 7.0) | 3.0 (1.0 - 4.0) | 1.0 (1.0 - 2.0) | 0.0 (0.0 - 1.0) |
| Best joint reconstruction | 3.0 (2.0 - 4.0) | 1.0 (1.0 - 2.0) | 0.0 (0.0 - 0.0) | 2.0 (1.0 - 2.0) | 5.0 (4.0 - 6.0) | 3.0 (2.0 - 3.0) | 1.0 (1.0 - 2.0) | 0.0 (0.0 - 1.0) |

**Abbreviations:** PFAV, putative function-altering variant

## Notes

### Competing Interest Statement

The authors have declared no competing interest.

## References

1. R. H. Ree, S. A. Smith, Maximum Likelihood Inference of Geographic Range Evolution by Dispersal, Local Extinction, and Cladogenesis. Systematic Biology 57, 4–14 (2008).

2. J. B. Joy, R. H. Liang, R. M. McCloskey, T. Nguyen, A. F. Poon, Ancestral reconstruction. PLoS computational biology 12, e1004763 (2016).

3. L. J. Revell, Ancestral state reconstruction of phenotypic characters. Evolutionary Biology 1–25 (2025).

4. B. S. W. Chang, K. Jönsson, M. A. Kazmi, M. J. Donoghue, T. P. Sakmar, Recreating a Functional Ancestral Archosaur Visual Pigment. Molecular Biology and Evolution 19, 1483–1489 (2002).

5. G. N. Eick, J. W. Thornton, Evolution of steroid receptors from an estrogen-sensitive ancestral receptor. Molecular and cellular endocrinology 334, 31–38 (2011).

6. R. Merkl, R. Sterner, Ancestral protein reconstruction: techniques and applications. Biological chemistry 397, 1–21 (2016).

7. W. J. Mellon, et al., Leveraging Comparative Phylogenetics for Evolutionary Medicine: Applications to Comparative Oncology. [Preprint] (2025). Available at: https://www.biorxiv.org/content/10.1101/2025.02.11.637459v1 [Accessed 16 July 2025].

8. Z. T. Compton, et al., Cancer Prevalence Across Vertebrates. Res Sq rs.3.rs-3117313 (2023). 10.21203/rs.3.rs-3117313/v1.

9. J. T. Kratzer, et al., Evolutionary history and metabolic insights of ancient mammalian uricases. Proceedings of the National Academy of Sciences 111, 3763–3768 (2014).

10. T. J. Barkman, Applications of ancestral sequence reconstruction for understanding the evolution of plant specialized metabolism. Philosophical Transactions of the Royal Society B: Biological Sciences 379, 20230348 (2024).

11. M. Bruto, et al., Ancestral gene acquisition as the key to virulence potential in environmental Vibrio populations. ISME J 12, 2954–2966 (2018).

12. M. A. Spyrou, K. I. Bos, A. Herbig, J. Krause, Ancient pathogen genomics as an emerging tool for infectious disease research. Nat Rev Genet 20, 323–340 (2019).

13. R. Agudo, M. P. Reche, Revealing antibiotic resistance’s ancient roots: insights from pristine ecosystems. Front. Microbiol. 15 (2024).

14. K. J. Gontjes, et al., Phylogenetic context of antibiotic resistance provides insights into the dynamics of resistance emergence and spread. J Infect Dis jiaf478 (2025). 10.1093/infdis/jiaf478.

15. X. Wang, et al., De novo acquisition of antibiotic resistance in six species of bacteria. Microbiol Spectr 13, e0178524 (2025).

16. K. C. Tracy, et al., Reversion to sensitivity explains limited transmission of resistance in a hospital pathogen. eLife 13 (2024).

17. Z. Yang, S. Kumar, M. Nei, A New Method of Inference of Ancestral Nucleotide and Amino Acid Sequences. Genetics 141, 1641–1650 (1995).

18. D. L. Swofford, W. P. Maddison, Reconstructing ancestral character states under Wagner parsimony. Mathematical Biosciences 87, 199–229 (1987).

19. M. Pagel, The Maximum Likelihood Approach to Reconstructing Ancestral Character States of Discrete Characters on Phylogenies. Systematic Biology 48, 612–622 (1999).

20. M. Pagel, A. Meade, D. Barker, Bayesian Estimation of Ancestral Character States on Phylogenies. Systematic Biology 53, 673–684 (2004).

21. D. Schluter, T. Price, A. Ø. Mooers, D. Ludwig, LIKELIHOOD OF ANCESTOR STATES IN ADAPTIVE RADIATION. Evolution 51, 1699–1711 (1997).

22. S. Ekman, H. L. Andersen, M. Wedin, The Limitations of Ancestral State Reconstruction and the Evolution of the Ascus in the Lecanorales (Lichenized Ascomycota). Systematic Biology 57, 141–156 (2008).

23. J. N. Keating, What is the best method for estimating ancestral states from discrete characters? [Preprint] (2023). Available at: https://www.biorxiv.org/content/10.1101/2023.08.31.555762v2 [Accessed 21 February 2025].

24. T. Pupko, I. Pe, R. Shamir, D. Graur, A Fast Algorithm for Joint Reconstruction of Ancestral Amino Acid Sequences. Molecular Biology and Evolution 17, 890–896 (2000).

25. J. D. Boyko, B. C. O’Meara, dentist: Quantifying uncertainty by sampling points around maximum likelihood estimates. Methods in Ecology and Evolution 15, 628–638 (2024).

26. M. Q. R. Pembury Smith, G. D. Ruxton, Effective use of the McNemar test. Behav Ecol Sociobiol 74, 133 (2020).

27. J. H. Han, et al., Whole-Genome Sequencing To Identify Drivers of Carbapenem-Resistant Klebsiella pneumoniae Transmission within and between Regional Long-Term Acute-Care Hospitals. Antimicrobial Agents and Chemotherapy 63, 10.1128/aac.01622-19 (2019).

28. L. Van der Maaten, G. Hinton, Visualizing data using t-SNE. Journal of machine learning research 9 (2008).

29. M. Ester, H.-P. Kriegel, J. Sander, X. Xu, Density-based spatial clustering of applications with noise in Int. Conf. Knowledge Discovery and Data Mining, (1996).

30. K. Saund, E. S. Snitkin, Hogwash: three methods for genome-wide association studies in bacteria. Microbial Genomics 6, e000469 (2020).

31. J. P. Allen, E. Snitkin, N. B. Pincus, A. R. Hauser, Forest and Trees: Exploring Bacterial Virulence with Genome-wide Association Studies and Machine Learning. Trends in Microbiology 29, 621–633 (2021).

32. S. J. Freeland, B. McCabe, Fitness compensation and the evolution of selfish cytoplasmic elements. Heredity 78, 391–402 (1997).

33. A. C. Palmer, et al., Delayed commitment to evolutionary fate in antibiotic resistance fitness landscapes. Nature communications 6, 7385 (2015).

34. H.-K. Kong, et al., Fine-tuning carbapenem resistance by reducing porin permeability of bacteria activated in the selection process of conjugation. Scientific reports 8, 15248 (2018).

35. F. Baquero, et al., Evolutionary pathways and trajectories in antibiotic resistance. Clinical Microbiology Reviews 34, e00050–19 (2021).

36. M. Pagel, Detecting correlated evolution on phylogenies: a general method for the comparative analysis of discrete characters. Proceedings of the Royal Society of London. Series B: Biological Sciences 255, 37–45 (1994).

37. M. Pagel, A. Meade, Bayesian Analysis of Correlated Evolution of Discrete Characters by Reversible-Jump Markov Chain Monte Carlo. The American Naturalist 167, 808–825 (2006).

38. J. D. Boyko, J. M. Beaulieu, Generalized hidden Markov models for phylogenetic comparative datasets. Methods in Ecology and Evolution 12, 468–478 (2021).

39. J. D. Boyko, B. C. O’Meara, J. M. Beaulieu, A novel method for jointly modeling the evolution of discrete and continuous traits. Evolution 77, 836–851 (2023).

40. J. D. Boyko, J. M. Beaulieu, Reducing the biases in false correlations between discrete characters. Systematic Biology 72, 476–488 (2023).

41. L. J. Revell, Ancestral state reconstruction of phenotypic characters. (2024).

42. G. James, D. Witten, T. Hastie, R. Tibshirani, An introduction to statistical learning: with applications in R (Springer, 2013).

43. M. Whitlock, D. Schluter, The analysis of biological data (Roberts Publishers Greenwood Village, Colorado, 2015).

44. S. J. Gould, Wonderful life: the Burgess Shale and the nature of history (WW Norton & Company, 1989).

45. B. C. O’Meara, Evolutionary Inferences from Phylogenies: A Review of Methods. Annual Review of Ecology, Evolution, and Systematics 43, 267–285 (2012).

46. V. Hanson-Smith, B. Kolaczkowski, J. W. Thornton, Robustness of Ancestral Sequence Reconstruction to Phylogenetic Uncertainty. Mol Biol Evol 27, 1988–1999 (2010).

47. M. N. Puttick, Partially incorrect fossil data augment analyses of discrete trait evolution in living species. Biology Letters 12, 20160392 (2016).

48. O. Gascuel, M. Steel, A Darwinian Uncertainty Principle. Syst Biol 69, 521–529 (2020).

49. R. Ponti, A. Arcones, D. R. Vieites, Challenges in estimating ancestral state reconstructions: the evolution of migration in Sylvia warblers as a study case. Integrative Zoology 15, 161–173 (2020).

50. T. A. Monson, et al., Keeping 21st Century Paleontology Grounded: Quantitative Genetic Analyses and Ancestral State Reconstruction Re-Emphasize the Essentiality of Fossils. Biology 11, 1218 (2022).

51. J. M. Beaulieu, B. C. O’Meara, Diversity and skepticism are vital for comparative biology. American Journal of Botany 106, 613–617 (2019).

52. H. Han, et al., Explainable t-SNE for single-cell RNA-seq data analysis. (2022). 10.1101/2022.01.12.476084.

53. J. Beaulieu, B. O’Meara, J. Oliver, J. Boyko, corHMM: Hidden Markov Models of Character Evolution. (2022).

54. M. W. Pennell, et al., geiger v2. 0: an expanded suite of methods for fitting macroevolutionary models to phylogenetic trees. Bioinformatics 30, 2216–2218 (2014).

55. B. R. Holland, S. Ketelaar-Jones, A. R. O’Mara, M. D. Woodhams, G. J. Jordan, Accuracy of ancestral state reconstruction for non-neutral traits. Scientific Reports 10, 7644 (2020).

56. J. D. Boyko, Automatic Discovery of Optimal Discrete Character Models. bioRxiv 2024–11 (2024).

57. H. L. Zhang, et al., Characterization of resistance to newer antimicrobials among carbapenem-resistant Klebsiella pneumoniae in the post–acute-care setting. Infection Control & Hospital Epidemiology 44, 1159–1162 (2023).

58. C. Collins, X. Didelot, A phylogenetic method to perform genome-wide association studies in microbes that accounts for population structure and recombination. PLOS Computational Biology 14, e1005958 (2018).

59. K. Saund, E. S. Snitkin, Hogwash: three methods for genome-wide association studies in bacteria. Microbial Genomics 6, e000469 (2020).

